# A Community-Developed Extension to Darwin Core for Reporting the Chronometric Age of Specimens

**DOI:** 10.1101/2021.11.24.469822

**Authors:** Laura Brenskelle, John Wieczorek, Edward Davis, Neill J. Wallis, Kitty Emery, Michelle J. LeFebvre, Rob Guralnick

## Abstract

Darwin Core, the data standard used for sharing modern biodiversity and paleodiversity occurrence records, has previously lacked proper mechanisms for reporting what is known about the estimated age range of specimens from deep time. This has led to data providers putting these data in fields where they cannot easily be found by users, which impedes the reuse and improvement of these data by other researchers. Here we describe the development of the Chronometric Age Extension to Darwin Core, a ratified, community-developed extension that enables the reporting of ages of specimens from deeper time and the evidence supporting these estimates. The extension standardizes reporting about the methods or assays used to determine an age and other critical information like uncertainty. It gives data providers flexibility about the level of detail reported, focusing on the minimum information needed for reuse while still allowing for significant detail if providers have it. Providing a standardized format for reporting these data will make them easier to find and search and enable researchers to pinpoint specimens of interest for data improvement or accumulate more data for broad temporal studies. The Chronometric Age Extension was also the first community-managed vocabulary to undergo the new Biodiversity Informatics Standards (TDWG) review and ratification process, thus providing a blueprint for future Darwin Core extension development.

## Introduction

Natural history specimens derived from paleontological or archaeological contexts (herein referred to as paleodiversity specimens for brevity) are critical for understanding past and present biodiversity and Earth history (Rick and Lockwood, 2013). The temporal contexts of paleodiversity specimens are essential to their value and use, and specific chronometric age detail is required for many analyses. Yet, open biodiversity repositories that aggregate digital data about these specimens, such as VertNet, iDigBio, and the Global Biodiversity Information Facility (GBIF; http://www.gbif.org), have had limited capabilities to provide information about absolute chronometric age. Previously, if this information was reported at all in biodiversity repositories, there were no best practices about what to report or where such reporting should be placed in standard fields used to support interoperability, greatly impeding the ability of users to find age information and complicating data reuse.

The Darwin Core (DwC) data standard, which is used for sharing bio- and paleodiversity specimen records, does have fields for geologicalContext (https://dwc.tdwg.org/terms/#geologicalcontext), which are inherently tied to chronology (Wieczorek et al., 2012), but there was no structure within this class by which chronometric age could be reported. Although geological context of especially paleontological specimens is critical information, and generally agreed-upon stratigraphic chronologies are published in the literature, the numerical age or age range associated with these contexts is not fixed, and thus may shift depending on current chronometric knowledge and the assigned chronology of an associated context. Because of this potential for shifting ages, the date when a specimen was assigned a specific geological context may impact whether or not this context still applies in the most current definition. Additionally, for archaeological specimens, geological context terms are insufficient because human-associated timelines are typically at a higher resolution, including decadal, centennial, or millennial timescales (LeFebvre et al., 2019). Paleontological specimens with ages that vary at those scales are also not sufficiently described using geologicalContext terms.

A key field for capturing temporal context in the current Darwin Core standard is eventDate. The definition of eventDate is “the date-time when the event was recorded” (https://dwc.tdwg.org/terms/#dwc:eventDate), and that definition explicitly also states that values for this term are “not suitable for a time in a geological context.” For specimens that are not from paleontological or archaeological contexts, this field is meant to capture when the specimen was collected, which is typically nearly contemporaneous with when the organism was last alive in its context. However, for occurrences from deeper time, it is unclear whether the ‘event’ is best applied to the collection date of a specimen or when this organism was last alive or deposited within the context from which it was recovered. Furthermore, this ambiguity of definition and application creates another source of confusion when deeper time occurrence records are intermixed with modern ones, which is the case in many biodiversity repositories. Finally, there previously were *no* dedicated fields in existing Darwin Core standards for describing what is known about the specific chronometric age of paleodiversity specimens (MacFadden and Guralnick, 2017). Taken together, these challenges in reporting chronometry limit the discoverability and use of these specimen records.

To better provide the necessary information about the chronometric age of paleodiversity specimens and to facilitate search functions by chronological age, we have developed a chronometric extension to the Darwin Core standard for reporting what is known about the absolute temporal context of the origin of specimens, and how this information is known (i.e. the evidence supporting an age). The Chronometric Age Extension (CAE; TDWG Darwin Core Chronometric Age Extension Task Group, 2021) is meant to support information about the multiple ways that chronometric ages are generated, whether from relative or absolute dating methods, age models that have been created for particular sites, or legacy collections information.

We place our development of the CAE in the larger context of standards development in the biodiversity community. Extensions such as CAE are not part of the Darwin Core standard; they are optional packages of terms that can be recorded to reflect additional information about biodiversity records and can reflect a many-to-one relationship of several extension datasets to a single specimen record (Droege et al., 2014; Endresen and Knüpfer, 2012). Extensions have served a valuable role for sharing extended specimen information, but have not been managed in the same way as core vocabularies, which go through a formal ratification process via standards organizations such as Biodiversity Informatics Standards (TDWG). As we discuss in detail below, there have been deficiencies with the way that extensions were built, managed, and deployed, and one aim of this paper is to provide an example of how extension development and ratification can be aligned with best practices used for core vocabularies and managed in the longer term.

Here, we first provide an empirical assessment of how chronometric age information is currently reported in global specimen occurrence publishing systems, considering published paleontological and archaeological resources. We use this assessment to highlight challenges and gaps with current practices, and then showcase the CAE to the Darwin Core, describing the goals of the extension, its current terms, and choices made in its development. We focus in particular on the stringent and long review and consultation that went into this effort, on the ratification process of the extension standard, and its management. We also present published examples of the CAE from the paleontological and archaeological communities and highlight how this effort aligns with others in reporting chronometric ages. Finally, we present a longer-term view of how to manage and grow new vocabularies more generally, based on lessons learned in developing the process for the CAE. Data standards like Darwin Core and its extensions are never fully “finished” products. They are meant to be maintained via community processes where feedback is welcomed and incorporated into the standard. We discuss how feedback during key phases in development was critical for the development of the CAE, in particular how to best balance broad applicability to all deeper-time specimens, while still serving needs for particular use cases.

## Materials and Methods

### Assessing gaps in current temporal reporting practices

We used a snapshot of data from GBIF taken on January 12, 2021 to understand current chronometric age reporting practices. In particular, we wanted to understand which Darwin Core fields might contain chronometric ages and what information is reported when chronometric age information is present. We downloaded all records labeled as a “FossilSpecimen” in dwc:basisOfRecord (zooarchaeological records are currently also labeled as “FossilSpecimen” in the DwC). To determine whether records contain explicit chronometric reporting, we targeted fields into which users were likely to input temporal information (Table 1). This predetermined list of search terms was developed via exploratory searches and in consultation with archaeologists and paleontologists. We aggregated unique values for the targeted Darwin Core fields and used search terms focused primarily on commonly used absolute time units. We used the grep() function in R to create a count of how many values were found to match these search terms in each field.

**Table 1.** A summary of the key fields and search terms for locating chronometry content. The first column shows the targeted fields within Darwin Core where chronometric data may be reported by providers. The second column provides a list of commonly used terms that are often used when reporting chronometry data.

| *FossilSpecimen Analysis* |  |
| --- | --- |
| Targeted Darwin Core Fields | Predetermined Search Terms |
| eventDate<br>verbatimEvent<br>dynamicProperties<br>identificationRemarks<br>lithostratigraphicTerms | "BCE", "BC"<br>"CE", "AD"<br>"BP"<br>"Ky"<br>"Kya"<br>"My"<br>"Ma"<br>"Mya"<br>"dating method" |

### Defining the goals of the extension

We intended for the extension to enable standard reporting of what is known about the chronometric age of a specimen, and if possible, how this information is known. This allows data users to assess fitness for use, provides them information on where to look (if anywhere) for more details about how the age was determined, and opens the door for reuse of these data in research. It also enables researchers to find specimens where age information could be improved with future assays.

Throughout the development of the CAE, we aimed to strike a balance between specific user needs and use cases and the following overarching goals to ensure the extension is usable for many different communities. These broad requirements are summarized succinctly below.

1. The extension, by definition, always links a specimen to a chronometric age context and is not meant to be deployed outside the context of that linkage.
2. The list of terms must be inclusive and support both relative and absolute dating approaches.
3. The list of terms must be able to capture date ranges and uncertainty based on different dating methods.
4. The list of terms must be able to support reporting on the assay or methods used to determine chronometric age.
5. The extension must be flexible to allow for various use cases across multiple disciplines (e.g. archaeology, paleontology).
6. The extension must not be too complex or convoluted for data providers to understand.

We kept these requirements in mind throughout the feedback and ratification processes and weighed community feedback against these as a way to test whether the extension was becoming too specific to a particular use case or discipline.

### Development of the Chronometric Extension

The results of the temporal reporting practices analysis (see below) suggest a strong need for a controlled vocabulary that can express not only chronometric ages associated with specimens, but also how such assays or determinations were performed and the degree of uncertainty in the chronometric data. To begin developing this vocabulary, we started with a set of four exemplar zooarchaeological datasets (Parnell, Tick Island, North Midden, and Baptizing Springs) published through VertNet as a part of the ZooArchNet project (LeFebvre et al., 2019). These datasets were purposefully picked for their differing types of chronometry reporting and are part of larger efforts towards mobilizing zooarchaeological specimen information at the Florida Museum of Natural History.

These use case datasets were critical for developing the Chronometric Age Extension because they provided challenging tests. As an example, the Parnell site assemblage (http://ipt.vertnet.org:8080/ipt/resource?r=flarch_zooarch_parnell_feature1) has both radiocarbon dating and ceramic seriation-based dating. The CAE had to be flexible enough to support reporting and uncertainty from both of these dating approaches. This flexibility is critical because the purpose of the extension is to allow data publishers to provide as deep or shallow structured chronometric age information associated with a specimen as they feel appropriate. The granularity of data reported in the extension should depend on a publisher’s views on usability or trustworthiness of the data. For example, a data publisher could choose to publish separate extension records that provide granular information about multiple assays used to assess absolute age context. Another, more conservative publisher might only publish a single, synthetic extension record with associated reporting on input source data and synthesis approach.

Based on these trials and extensive discussion, especially with practicing archaeologists, we developed an initial set of extension terms and definitions. We developed the initial list of terms as a minimum set of fields to enable the assessment of quality and usability of the data with a focus on the methods used to generate absolute age context, however generated. We then assessed the utility of these terms to describe absolute chronometric context for specimen data from the four exemplar zooarchaeological datasets. Co-author Wallis provided the Beta Analytic output report for the Parnell radiocarbon dates to illustrate the information returned to investigators who have these analyses performed. Using input from the archaeologists on the team, we revised the initial term list in a loop of iterative feedback, editing the extension draft until we had developed terms and definitions on which we all agreed.

Once we reached an internal consensus about these terms, their labels, and definitions within the smaller, initial development group based on the archaeological perspective, we worked with paleontological collaborators utilizing test datasets (see Tables S6-S8) to determine whether the extension would be usable for reporting information about chronometric ages related to paleontological specimens. We used feedback from this process to further improve our term list. Next, we reached out to other experts in the paleontology and archaeology communities to review how the extension worked in practice. Their input helped call attention to terms that were too specific to our first test cases or definitions that were unclear to users. The lead author also presented at a Darwin Core Hour, which is a community forum for presenting about topics related to natural history collections standards, on April 24, 2018 (available at https://github.com/tdwg/dwc-qa/wiki/Webinars#chapter13). We used this forum to solicit feedback from the broader community and explain more about the intent and utility of the extension effort. Subsequently, the lead author, together with co-authors Guralnick and Wieczorek, worked with TDWG to express the extension formally, following the XML schema-based model for Darwin Core extensions (https://github.com/gbif/rs.gbif.org/blob/master/schema/extension.xsd).

### Testing the Chronometric Age extension

We tested the ability of the extension to meet the needs of data providers by publishing specimen data from zooarchaeological and paleontological collections to VertNet. This process began in 2018 with the prototype of the extension and continued until the final ratification of the extension by TDWG (see below). First, we assessed whether the extension allowed sharing of all data that domain scientists felt was necessary for reuse and quality assessment. We also evaluated whether the extension field names and definitions were clearly understandable by users who did not participate in extension development. This was an iterative process of helping data providers find the appropriate fields for different pieces of information if necessary, then discussing with them whether or not these fields and values made sense for the data they were trying to share. We also iteratively discussed whether the extension fields covered all concepts necessary to report their data with collaborating data providers and domain scientists. Throughout this feedback process, we made changes based on user suggestions to improve clarity and better suit user needs.

### Chronometric Age extension formal ratification process

In April 2020, we officially created a Task Group under the TDWG Earth Sciences and Paleontology Interest Group (https://www.tdwg.org/community/esp/chrono/) to develop and ratify the Chronometric Age extension to Darwin Core as an official TDWG-approved data standard for biodiversity data. This task group helped formalize term definitions and test the proposed extension further with the publication of new datasets. Following TDWG Vocabulary Maintenance Standard protocol (https://www.tdwg.org/standards/vms/), the extension was opened for a formal public review in November 2020. This review was advertised through various listservs to communities with a possible vested interest in the extension. To ensure this review was fully transparent and documented, the process was done on GitHub so anyone could comment with any questions, concerns, or proposed changes (https://github.com/tdwg/chrono/issues/15). The review was initially for a 30-day period. If a reviewer’s comment led to a consensus agreement that required a change or addition to the extension, that change was made and the review was extended for an additional 30 days. This resulted in three rounds of changes to the extension, and the review officially ended in March 2021.

Once the review was complete, the extension was submitted for formal approval as a vocabulary enhancement to Darwin Core by the TDWG Technical Architecture Group and Executive Committee. As part of submitting the Chronometric Age extension for approval by TDWG, we included equivalent terms found in ABCD EFG (Access to Biological Collections Data Extended For Geoscience). ADBC EFG is another TDWG-approved data standard for geoscience collections that is commonly used but whose remit is broader and less focused on minimum reporting requirements. These equivalences can be found in the current CAE terms list (https://github.com/tdwg/chrono/blob/master/vocabulary/term_versions.csv) under the heading ‘abcd_equivalence’. In April 2021, the Chronometric Age extension was ratified as part of Darwin Core by TDWG. We then finalized the schema for the extension to be compatible with publishing via Darwin Core archive.

## Results

### Results of Current Temporal Reporting Practices Analysis

We assembled all the records with basisOfRecord recorded as ‘FossilSpecimen’ and evaluated unique value counts and chronometry-related values (see Table 1). Reporting of chronometric information is sparse and scattered across multiple Darwin Core fields. Table 2 shows the number of unique values and the number of chronometry-related values we found for each of the targeted Darwin Core fields in our dataset. The reporting in these fields is highly variable. In many cases, chronometric information is bundled with other information and placed in unexpected Darwin Core fields where it would be hard for a user to find it.

**Table 2.** Summary analysis of the records with basisOfRecord recorded as FossilSpecimens, with the number of unique values found in each targeted Darwin Core field and the number of chronometry-related values found within each field.

| Darwin Core Field | Unique Value Counts | Chronometry-Related Values |
| --- | --- | --- |
| eventDate | 6497 | 0 |
| verbatimEvent | 14564 | 0 |
| dynamicProperties | 358284 | 144 |
| identificationRemarks | 49523 | 32 |
| lithostratigraphicTerms | 85013 | 3153 |

### Unveiling the Ratified Chronometric Extension

The terms in the CAE were defined following the model used for the Darwin Core standard. This model consists of formal definitions of the terms and their complete history captured in a CSV file with a row for each version of a term and fields that describe the properties of each term version, including, among other attributes, an identifier, preferred label, definition, examples, and whether a version is recommended for current use. The current version of this file can be found at https://github.com/tdwg/chrono/blob/master/vocabulary/term_versions.csv. Deprecated terms from the long, iterative development process can be found at the bottom of the CSV file. A compressed version is shown in Table 3.

**Table 3.** The current Chronometric Age extension terms and definitions with notes detailing examples of what is meant by specific field names. This same information, including notes detailing examples of what is meant by each specific field, can be found at the current published version of the extension on the TDWG website at http://rs.tdwg.org/chrono/version/terms/2021-02-21.htm.

| Term | Definition |
| --- | --- |
| chronometricAge | An approximation of a temporal position (in the sense conveyed by https://www.w3.org/TR/owl-time/#time:TemporalPosition) that is supported via evidence. |
| chronometricAgeID | An identifier for the set of information associated with a ChronometricAge. |
| chronometricAgeReferences | A list (concatenated and separated) of identifiers (publication, bibliographic reference, global unique identifier, URI) of literature associated with the ChronometricAge. |
| chronometricAgeRemarks | Notes or comments about the ChronometricAge. |
| materialDated | A description of the material on which the chronometricAgeProtocol was actually performed, if known. |
| materialDatedID | An identifier for the MaterialSample on which the chronometricAgeProtocol was performed, if applicable. |
| materialDatedRelationship | The relationship of the materialDated to the subject of the ChronometricAge record, from which the ChronometricAge of the subject is inferred. |
| chronometricAgeProtocol | A description of or reference to the methods used to determine the chronometric age. |
| uncalibratedChronometricAge | The output of a dating assay before it is calibrated into an age using a specific conversion protocol. |
| verbatimChronometricAge | The verbatim age for a specimen, whether reported by a dating assay, associated references, or legacy information. |
| earliestChronometricAge | The maximum/earliest/oldest possible age of a specimen as determined by a dating method. |
| earliestChronometricAgeReferenceSystem | The reference system associated with the earliestChronometricAge. |
| latestChronometricAge | The minimum/latest/youngest possible age of a specimen as determined by a dating method. |
| latestChronometricAgeReferenceSystem | The reference system associated with the latestChronometricAge. |
| chronometricAgeConversionProtocol | The method used for converting the uncalibratedChronometricAge into a chronometric age in years, as captured in the earliestChronometricAge, earliestChronometricAgeReferenceSystem, latestChronometricAge, and latestChronometricAgeReferenceSystem fields. |
| chronometricAgeUncertaintyInYears | The temporal uncertainty of the earliestChronometricAge and latestChronometricAge in years. |
| chronometricAgeUncertaintyMethod | The method used to generate the value of chronometricAgeUncertaintyInYears. |
| chronometricAgeDeterminedBy | A list (concatenated and separated) of names of people, groups, or organizations who determined the ChronometricAge. |
| chronometricAgeDeterminedDate | The date on which the ChronometricAge was determined. |

Figure 1 shows the conceptual basis for the extension, with terms expressing metadata about the whole chronometric dating process, capturing information about the material dated to determine the age, the dating analysis itself, and the age inference of the specimen based on the dating analysis. The color-coding in this figure has been carried through to Table 3, which shows the terms and definitions of the current, ratified Chronometric Age extension. This same list of terms with comments, examples, and other information can be found at https://github.com/tdwg/chrono/blob/master/vocabulary/term_versions.csv.

**Figure 1.**
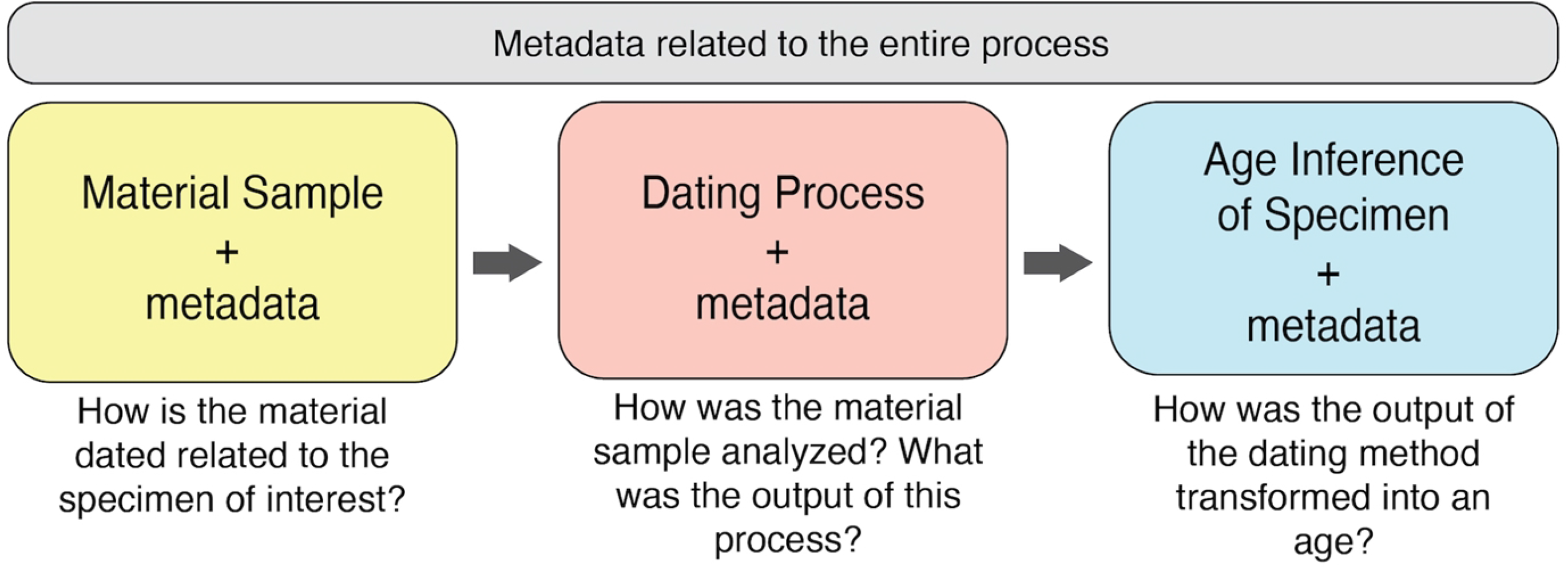
The conceptual layout of the Chronometric Age extension with data representing different parts of the dating process.

### Exemplar Use Cases

To test the usability of the Chronometric Age Extension during development, we published paleontological and archaeological datasets. This process allowed us to refine the list of necessary terms for a wide array of use cases and to clarify definitions that users found ambiguous or hard to understand. Here we have highlighted three published specimen records from three separate datasets that were dated using different methods to show the breadth and flexibility of the extension.

#### North Midden

North Midden (http://ipt.vertnet.org:8080/ipt/resource?r=north_midden) is an archaeological site located in Florida, U.S.A. The age for the zooarchaeological specimens from the site is based on an AMS analysis performed on the right valve of a *Crassostrea virginica* specimen found in Feature 17 at the site. Figure 2 shows the conceptualization for this example in a published Chronometric Age extension, and Table S2 shows a published example record.

**Figure 2.**
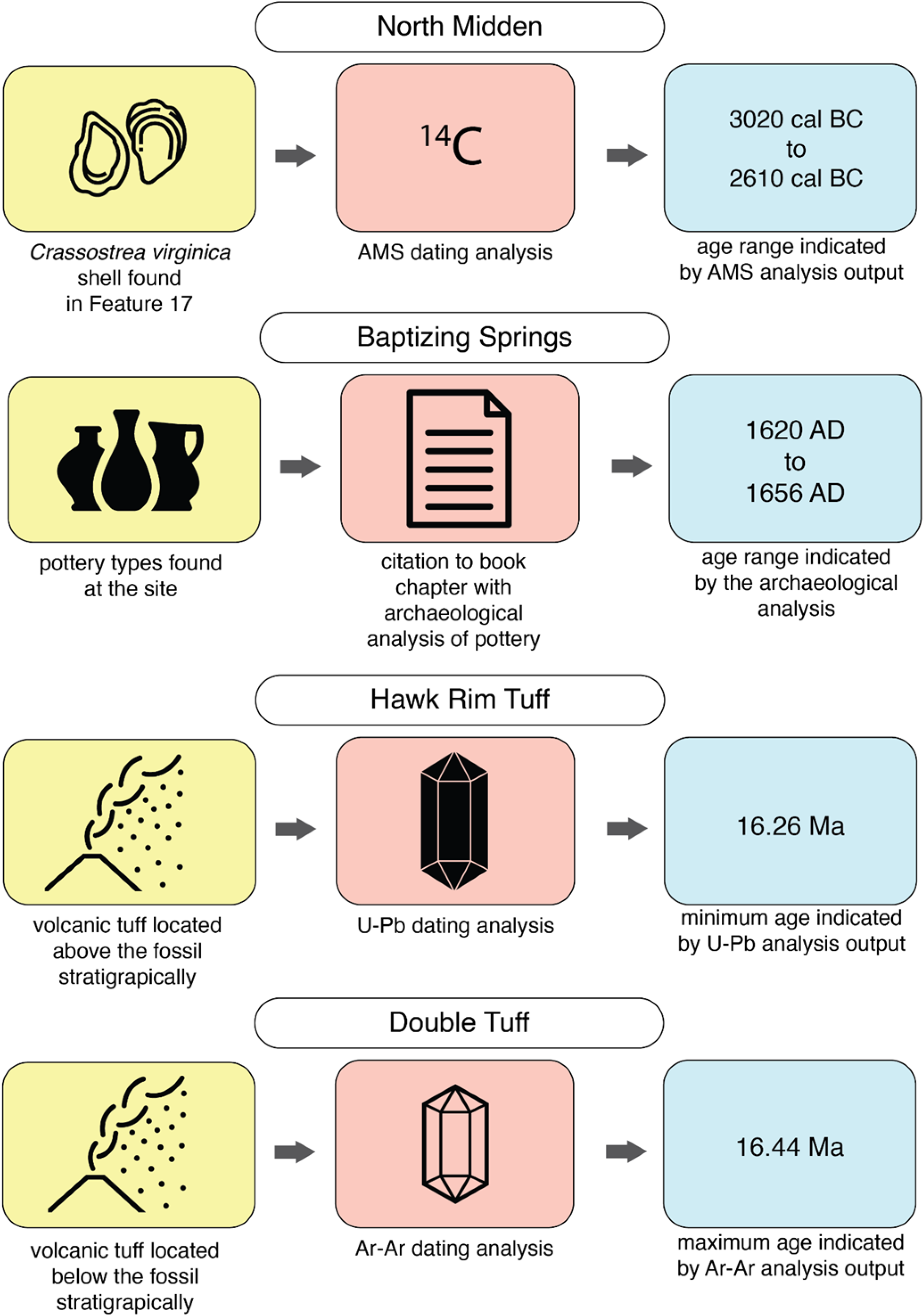
Exemplar material samples, data assays and outcomes of the three example datasets, with color coding to indicate how the different parts of the dating process correspond with the CAE terms in Table 3 and conceptual elements in Figure 1. “Oyster” icon by Eucalyp, “Ceramics” icon by Adrien Coquet, “Article” icon created by IconMark, “Volcanic Ash” icon created by Hayashi Fumihiro, “Crystal” icons created by MarkieAnn Packer all from http://www.thenounproject.com.

#### Baptizing Springs

Baptizing Springs (http://ipt.vertnet.org:8080/ipt/resource?r=baptizing_springs) is an archaeological site located in Florida, U.S.A. that we chose to use as another example use case because the age of the specimens from this site are based on a combination of historical documentation of Spanish mission sites in Florida and Spanish ceramics and other artifact types found at the site. The age concluded by the synthesis of these lines of evidence is reported in the extension record (Figure 2), and the details of these analyses are referenced in the chronometricAgeReferences field (Table S4).

#### Hawk Rim

The third example use case is from a paleontological site called Hawk Rim in Oregon, U.S.A. This use case was chosen because it highlights the flexibility of the extension. In Darwin Core, a specimen record can have multiple associated extension records. This enables reporting for numerous different chronometric dating assays, or in the case of Hawk Rim, separate extension records for reporting the earliest and latest possible ages in an age range, as indicated by two different dating assays on two different material samples (Figure 2).

## Discussion and Future Directions

### Current limitations with reporting deeper time and the value of a Chronometric Age extension

Darwin Core did not have a consistent, designated place to record specimen absolute age information. Because of this, data about a specimen’s absolute age have previously been reported in a variety of different Darwin Core fields, which makes them difficult for users to find and effectively use. In our empirical assessment of how deeper-time chronometric age is currently reported, we located information about absolute age not only in Geological Context fields such as lithostratigraphicTerms, but also in identificationRemarks and dynamicProperties (Table 2). Overall, absolute age information remains hidden across various fields even in standardized, published digital records of paleontological and archaeological specimens. This highlights the value of an extension to coherently report this information. The CAE is complementary and non-duplicative with any other dataset-level reporting of temporal scope. Published datasets can and do have their own metadata, but these are aggregations at the dataset-level and do not refer directly to specimens.

Because data that are stored in Darwin Core extensions are separate from the data in the core occurrence file of a Darwin Core Archive, they are not usually displayed in data aggregator portals (such as iDigBio or VertNet). In fact, to view extension data, a user must download a compressed Darwin Core Archive folder and view the extension file separately from the main occurrence data. We also note that there is no *a priori* reason why the same reporting cannot also be placed in dwc:dynamicProperties in a standard syntax (e.g. JSON such as in our exemplar datasets; Tables S3, S5, S8) to enhance discoverability. For this reason, in our example zooarchaeological datasets, we also published CAE data in a machine readable JSON format in dwc:dynamicProperties. This practice makes these data viewable on the aggregator portals such as GBIF and iDigBio and ensures that CAE data are found in the occurrence CSV file, which most users use to retrieve Darwin Core data.

One place we found no absolute age information reported was in the Darwin Core eventDate field (Table 2). The definition of that term is unclear whether the “event” to be reported is when the organism was alive in its context or when the specimen was collected. This confusion is based on Darwin Core “events” not explicitly being defined as sampling events in the standard. While the term definition may be ambiguous, data providers all seem to interpret this field to refer to the calendar date when a sampling or collection event happened, whether it is for a paleontological sample or neontological one. Given both current practice and the obvious need to report collection event dates, we recommend that eventDate continue to be used to refer to the “collection event” and the chronometry extension serve as the key mechanism for reporting the absolute age of a specimen. We also recommend a change to the definition of eventDate to more explicitly state that this is the appropriate place to record the date of collection for all specimen types.

### What the extension is and is not

During the review and community feedback process, we were asked about the ability of the extension to serve information about complex age models. The extension is capable of reporting the overall conclusions of an age model, and it is capable of reporting details of an age model to the extent that a data provider wants to include separate extension records for different pieces of evidence or information used to develop the age model. However, it was not our intention to develop an extension for reporting complex age models. Given the current state of reporting chronometric information for specimens in Darwin Core with the basisOfRecord ‘FossilSpecimen’, our main goal was to provide a format for publishing these data, in relation to specimen records, in a way that makes them more readily sharable and accessible. Because the extension is meant to serve many diverse use cases and disciplines, we felt it was important to keep the extension as simple as possible while also providing the necessary fields for reporting critical information for understanding the chronometric context of a specimen. It is important to note here that, as with all of Darwin Core, no fields are required to be filled in the extension. The three exemplar use cases provided in the Results demonstrate this. Similarly, as illustrated by the paleontological examples detailed above (Table 6 and Table 7), one specimen can have multiple associated Chronometric Age extension records, when necessary.

### Integrating information about absolute ages across repositories - next steps

The flexibility of the Chronometric Age extension facilitates the possibility of crosswalking data from other sources, such as Open Context (Kansa et al., 2014), Neotoma (Williams et al., 2018) or the Paleobiology Database (Peters and McClennen, 2015), in the future. As part of our current work, we have published chronometric data for example zooarchaeological and paleontological datasets, but it is important that we continue to work on publishing datasets with chronometric information from different communities that use assays or evidence types not represented by our examples, or in the use cases provided by community feedback. These efforts would ensure that the Chronometric Age extension does indeed work for all paleontological and archaeological specimens, and allow us to troubleshoot unforeseen problems or ambiguity in the terms or their definitions. Additionally, publishing more datasets with the CAE would expand the amount of findable paleontological and zooarchaeological data with absolute ages, further enabling research using occurrence records from particular temporal contexts.

The Chronometric Age extension also highlights an intersection of the disciplines of biodiversity and geological informatics. For example, the materialDated and materialDatedID terms can and often do refer to non-biological specimen materials. The identifiers for these materials can include International Geo Sample Numbers (IGSN; http://igsn.github.io/) and could link to digital repositories for geological sample information, such as the System for Earth Sample Registration (SESAR; http://www.geosamples.org/). With these methods, we hope to begin to bridge the gap between efforts in the biological and archaeological communities with those in geoinformatics.

### Building better extensions and the value of managed vocabularies

The Chronometric Age Extension was the first Darwin Core extension to undergo an official review and ratification process by TDWG. We provide here some lessons learned and best practices that emerged during that process. First, while the process of ratifying a standard requires considerably more time and effort than not doing so, it has tangible benefits. We received important feedback and questions from community members that made us reassess key definitions to make them less ambiguous. The open review process and use of the GitHub platform allowed us to engage with people beyond the realm of natural history museum informatics, and specific feedback we received from stakeholders in geoinformatics helped us to better define what we mean by “chronometric age” (https://github.com/tdwg/chrono/issues/27). Using GitHub also enabled us to document and archive the conversations that were held during the public review so the thought process behind changes and additions can be better understood by future contributors and users. All of the discussions and changes resulting from the review can be viewed on GitHub (https://github.com/tdwg/chrono/milestone/5?closed=1). Although this project began with a small group of collaborators to develop an initial set of terms and definitions, the ratification process helped broaden our reach significantly. In sum, the community feedback and TDWG ratification process were critical for improving the usability and clarity of the extension.

This ratification process within a standards organization provides a means for long-term management and sustainability of efforts because ratified standards become community owned, and that open process does not stop once ratification happens. Community data standards, while seeking long-term stability, are still open to further development. The Chronometric Age Extension GitHub repository (https://github.com/tdwg/chrono) persists and is open for comments and revisions as the community sees fit. An additional lesson learned is that it is also necessary to balance community feedback with the goals of a data standard to avoid overspecificity supporting only particular use cases. It is challenging to balance these two sometimes opposing approaches, but when done properly, it results in a flexible, functional data standard that can be used by many different disciplines. It is our hope that the Chronometric Age Extension will promote data discoverability for the ages of specimens, thus expanding and enabling research across broader temporal scales and across the disciplines of paleontology, archaeology, and geology.

## Acknowledgments

We would like to acknowledge the helpful insight provided by people who reviewed the extension during this process, including but not limited to Arianne Boileau, Phil Buckland, Nicole Cannarozzi, Scott Fitzpatrick, Christina Giovas, Carla Hadden, Samantha Hopkins, Jessica King, Holly Little, Scott Macrae, Amanda Millhouse, Paul Morris, Denné Reed, Stephen Richard, Ashley Sharpe, Vicki Szabo, and Victor Thompson.

## Notes

### Competing Interest Statement

The authors have declared no competing interest.

